# Seabird nutrient subsidies enrich mangrove ecosystems and are exported to nearby coastal habitats

**DOI:** 10.1101/2023.11.19.567759

**Authors:** Jennifer Appoo, Nancy Bunbury, Sébastien Jaquemet, Nicholas A.J. Graham

## Abstract

Eutrophication by human-derived nutrient enrichment is a major threat to mangroves, impacting productivity, ecological functions, resilience and ecosystem services. Natural mangrove nutrient enrichment processes, however, remain largely uninvestigated. Mobile consumers such as seabirds are important vectors of cross-ecosystem nutrient subsidies to islands but how they influence mangrove ecosystems is poorly known. We assessed the contribution, uptake, cycling and transfer of nutrients from seabird colonies in remote mangrove systems free of human stressors. We found that nutrients from seabird guano enrich mangrove plants, reduce nutrient limitations, enhance mangrove invertebrate food webs and are exported to nearby coastal habitats through tidal flow. We show that seabird nutrient subsidies in mangroves can be substantial, improving the nutrient status and health of mangroves and adjacent coastal habitats. Conserving mobile consumers, such as seabirds, is therefore vital to preserve and enhance their role in mangrove productivity, resilience and provision of diverse functions and services.

## INTRODUCTION

Mangrove forests occupies the interface between land and sea, where it plays critical roles in sustaining biodiversity and maintaining ecological functions and services ^1,2^. For example, mangrove habitat provides protection against coastal erosion and tidal surges ^3^; provides essential food, protection and habitat to numerous taxa during part or all of their life cycles ^4^; and sequesters large amounts of carbon, contributing substantially to climate change adaptation and mitigation ^5^. The provisioning of these functions and services are dependent on the status and health of mangrove forests ^6^, which are increasingly threatened by human activities. Eutrophication, caused by excessive human-derived nutrient inputs is a major threat for coastal ecosystems, including mangroves ^7^. Nutrient inputs from anthropogenic activities tend to be rich in nitrogen (N) but poor in phosphorus (P), and can reduce mangrove growth, cause death of pneumatophores and soil anoxia ^8^, and increase mortality of mangrove forests ^9^. Although strong links have been documented between human-derived nutrient enrichment and mangrove ecosystem functioning, there has been far less attention on the impacts of mangrove nutrient enrichment by natural sources.

Mobile consumers play a key role in the movement of nutrients across ecosystem boundaries^10^. Seabirds comprise an important group of mobile consumers involved in the transport of nutrients from sea to land, by feeding in oceanic areas and depositing large amounts of guano in their colonies^11^. Enriched in both N and P compounds, seabird-derived nutrients enhance primary productivity around their colonies, and in coastal habitats such as coral reefs, resulting in increased growth rate of reef fish ^12,13^ and coral ^14^, as well as increased fish biomass ^12^. Despite the widespread occurrence of seabirds nesting in mangroves ^7^, much less is known about the influence of seabird-derived nutrients in mangrove ecosystems. Previous studies documented positive relationships between seabird nutrient subsidies and mangrove productivity ^15^ and nutrient status ^16–18^, but these studies only focused on mangroves in northern and central America in proximity to urban centers. How seabird-derived nutrients influence mangrove forests in regions such as the extensive and mangrove-rich Indo-Pacific, and in the absence of human influence, is unknown. Additionally, no studies have explored the vertical transfer of seabird-derived nutrients within mangrove food webs. Furthermore, tides strongly mediate connectivity of mangroves with adjacent coastal habitats through the exchange of detritus, fauna and nutrients ^19^. Consequently, seabird influence may extend beyond mangroves in the coastal seascape; however, the horizontal transfer of seabird-derived nutrients across mangrove boundaries has not been studied. Documenting these relationships and linkages is increasingly important given the accelerated declines in both mangrove habitats ^20^ and seabird populations ^21,22^ due to numerous anthropogenic threats.

Here, we examined the contribution, uptake and transfer of seabird-derived nutrients in mangrove habitats on Aldabra Atoll, in the Southern Seychelles, one of the largest mangrove-nesting seabird colonies in the Indian Ocean. Aldabra hosts the largest area of mangroves in the Seychelles archipelago, which supports the largest breeding populations of frigatebirds, *Fregata minor* and *F. ariel*, in the Indian Ocean ^23^, as well as one of the largest breeding populations of red-footed boobies *Sula sula* in the region. Frigatebirds and red-footed boobies nest exclusively in mangrove forests around Aldabra’s lagoon shores ^24^. With only a small research station, human impact on Aldabra is minimal, presenting ideal conditions to assess the effects of seabird subsidies in a relatively undisturbed mangrove system. We aimed to answer the following questions: (1) What quantities of marine-derived nutrients are contributed by mangrove-nesting seabirds? (2) Are seabird-derived nutrients transferred to and assimilated by mangroves? (3) Do seabird-derived nutrients alleviate nutrient limitation and reduce nutrient resorption efficiency of mangroves? (4) Are seabird-derived nutrients trophically transferred to mangrove fauna? And (5) Are seabird-derived nutrients in mangroves exported to adjacent coastal habitats?

We achieved this using biogeochemical assays of multiple ecosystem components, including seabird guano, mangrove sediment, leaves, gastropods, crabs, adjacent macroalgae and seawater, sampled at 10 sites with (n = 5) and without (n = 5) nesting seabirds. Using seabird demography metrics, we find that seabird nutrient subsidies in mangroves can be substantial. Nutrient assays reveal seabird-derived nutrients enrich mangroves and reduce their nutrient limitations, but do not influence their resorption efficiency. Seabird-derived nutrients enrich mangrove invertebrates through trophic transfer and are exported to nearby coastal habitats through tidal flow. By showing that seabird-derived nutrients promote nutrient status and health of mangroves, our results increase understanding about the implications of declines of seabird populations, and suggest that efforts to maintain or restore their populations and breeding grounds should be prioritized.

## RESULTS

### Seabird nutrient contributions to mangroves are substantial

Red-footed booby guano contained 10.1 ± 2.92 %N and 10.2 ± 0.87 %P (Mean ± SD, n = 4) and frigatebird guano had 10.9 ± 2.06 %N and 10.3 ± 1.14 %P (Mean ± SD, n = 4). Guano nitrogen isotopic values (δ^15^N) for red-footed booby were 10.8 ± 2.71 (Mean ± SD, n = 20) and for frigatebirds 12.9 ± 3.63 (Mean ± SD, n = 20). Based on seabird population size and breeding metrics, and guano nutrient concentrations (see experimental procedures), estimated total annual nutrient contributions were higher for red-footed boobies (28.6 N tonne.year^−1^; 28.9 P tonne.year^−1^) than for frigatebirds (12.8 N tonne.year^−1^; 12.0 P tonne.year^−1^).

### Seabird-derived nutrients are transferred and assimilated by mangroves

We recorded mangrove forest metrics at all sites, resulting in a total of 923 surveyed trees, with *R. mucronata* making up 79% of individuals and being the dominant species at all surveyed sites (Table S1). δ^15^N in *R. mucronata* leaves and mangrove sediment were higher at seabird sites compared to non-seabird sites (Figure 1D, 1E; Table S2). Nutrient levels in *R. mucronata* leaves were higher in the presence of nesting seabirds (Table S2), for both N (Mean ± SD: seabird = 1.02 ± 0.21%, non-seabird = 0.73 ± 0.09%, *P* < 0.0001) and P (seabird = 0.09 ± 0.02%, non-seabird = 0.07 ± 0.01%, *P* = 0.01; Figure 2A, B).

**Figure 1.**
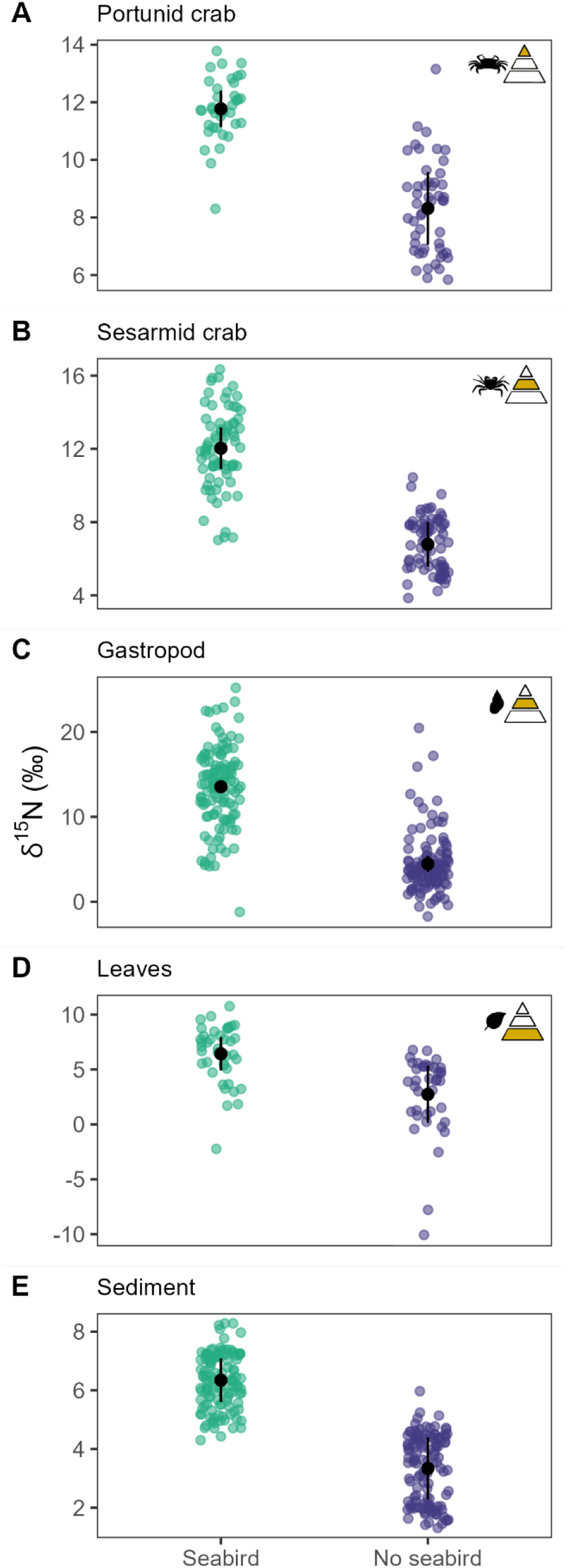
Nitrogen isotopic values in mangrove abiotic and biotic (trophic) components at seabird and non-seabird sites. δ^15^N values of portunid crabs *Thalamita crenata* (A), sesarmid crabs *Sesarma leptosoma* (B), gastropods *Littoraria* spp. (C), mangrove leaves *Rhizophora mucronata* (D) and mangrove sediment (E) at seabird and non-seabird breeding sites on Aldabra Atoll. Black points and error bars display predicted means ± SD of linear mixed models and green and purple points display raw data. Error bars not visible in some cases because of scaling. Trophic level of each biotic component is indicated by the triangular icon in the top right of each plot.

**Figure 2.**
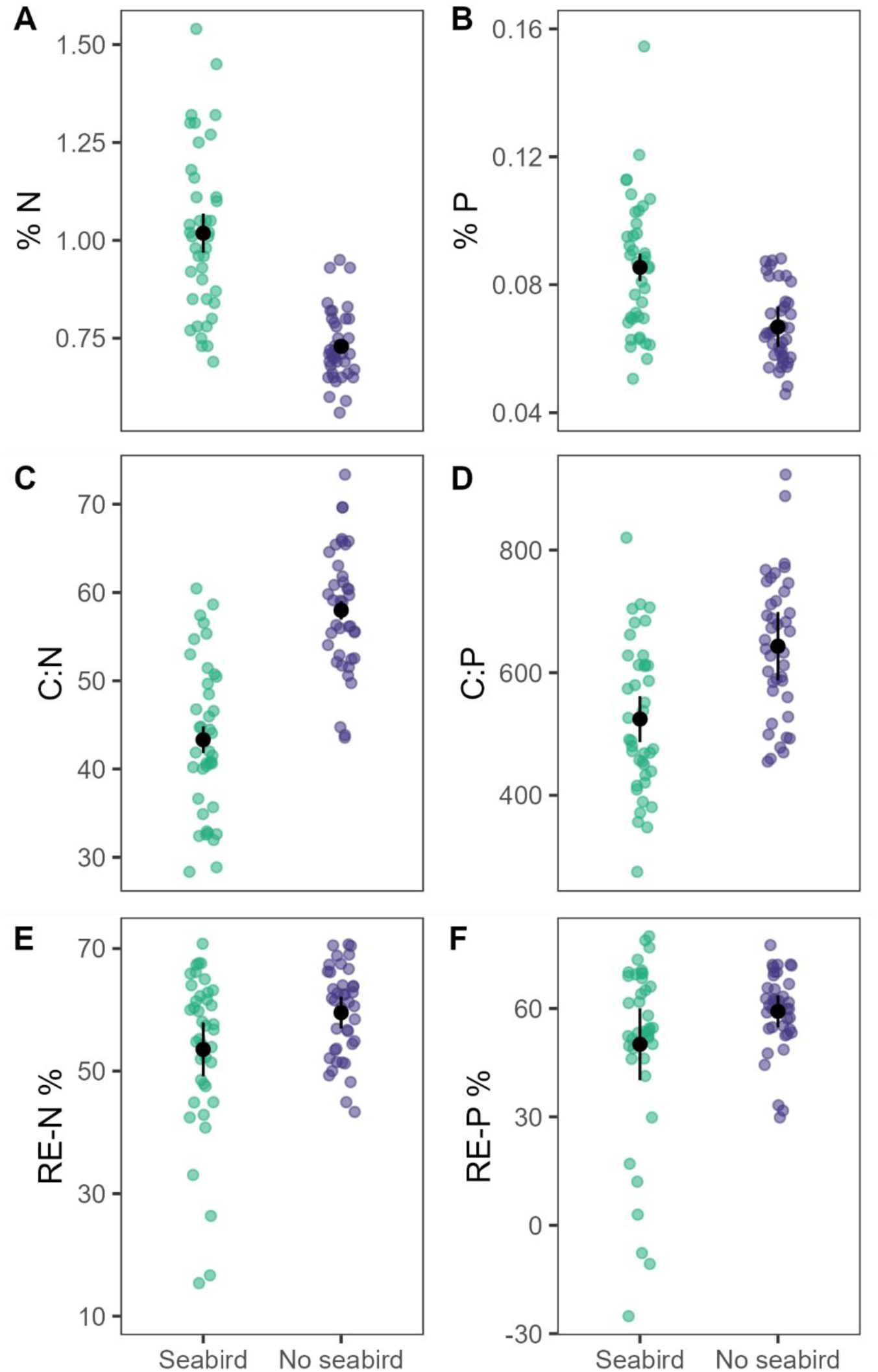
Nutrient parameters in mangrove leaves at seabird and non-seabird sites. Nutrient concentrations, %N (A) and %P (B), nutrient ratios C:N (C) and C:P (D), and nutrient resorption efficiencies RE-N (E) and RE-P (F) of mangrove leaves *Rhizophora mucronata* at seabird and non-seabird breeding sites on Aldabra Atoll. Black points and error bars display predicted means ± SD of linear mixed models and green and purple points display raw data. Error bars not visible in some cases because of scaling. RE-N: nitrogen resorption efficiency, RE-P: phosphorus resorption efficiency.

### Seabird-derived nutrients reduce mangrove nutrient limitations but not resorption efficiency

Nutrient ratios, an indicator for nutrient limitation, were lower at seabird sites (C:N: seabird = 43.3 ± 8.64, non-seabird = 58.0 ± 6.95, *P* < 0.0001; and C:P: seabird = 524.4 ± 122.6, non-seabird = 643.2 ± 115.0, *P* = 0.031; Figure 2C, D). The overall average N:P ratio of mangroves was 11.7 ± 2.45. We estimated mangrove nutrient resorption efficiency as an indicator for mangrove nutrient cycling, using nutrient concentrations of green and senescent leaves (see experimental procedures). Nutrient resorption efficiency was similar between sites with and without seabirds (N resorption efficiency [RE-N]: seabird = 53.5 ± 13.1%, non-seabird = 59.5 ± 7.39%, *P* = 0.15; and P resorption efficiency [RE-P]: seabird = 50.1 ± 25.2%, non-seabird = 59.2 ± 10.9%, *P* = 0.24; Figure 2E, F; Table S2).

### Seabird-derived nutrients are transferred to mangrove invertebrate food web

δ^15^N were higher at seabird sites compared to non-seabird sites at all trophic levels, including for herbivorous gastropods (seabird = 13.5 ± 4.64%, non-seabird = 4.45 ± 3.42%, *P* < 0.0001; Figure 1C), herbivorous sesarmid crabs (seabird = 12.03 ± 2.16%, non-seabird = 6.79 ± 1.47%, *P* < 0.001; Figure 1B) and omnivorous portunid crabs (seabird = 11.78 ± 1.09%, non-seabird = 8.32 ± 1.62%, *P* < 0.005; Figure 1A).

### Seabird-derived nutrients are exported to nearby coastal habitats

Surface seawater in the lagoon had higher NOx (nitrate + nitrite) at seabird sites than non-seabird sites only during outgoing tides (post-hoc tests: *P* = 0.033; Figure 3A; Table S3), whereas phosphate concentrations were higher at seabird than non-seabird sites during both incoming and outgoing tides (post-hoc tests: incoming: *P* < 0.0001, outgoing: *P* = 0.0051; Figure 3B; Table S3). Macroalgae *Halimeda* spp. sampled on substrates adjacent to mangroves had higher δ^15^N at seabird than non-seabird sites (seabird = 9.63 ± 1.62, non-seabird = 5.98 ± 0.91, *P* = 0.021; Figure 3C).

**Figure 3.**
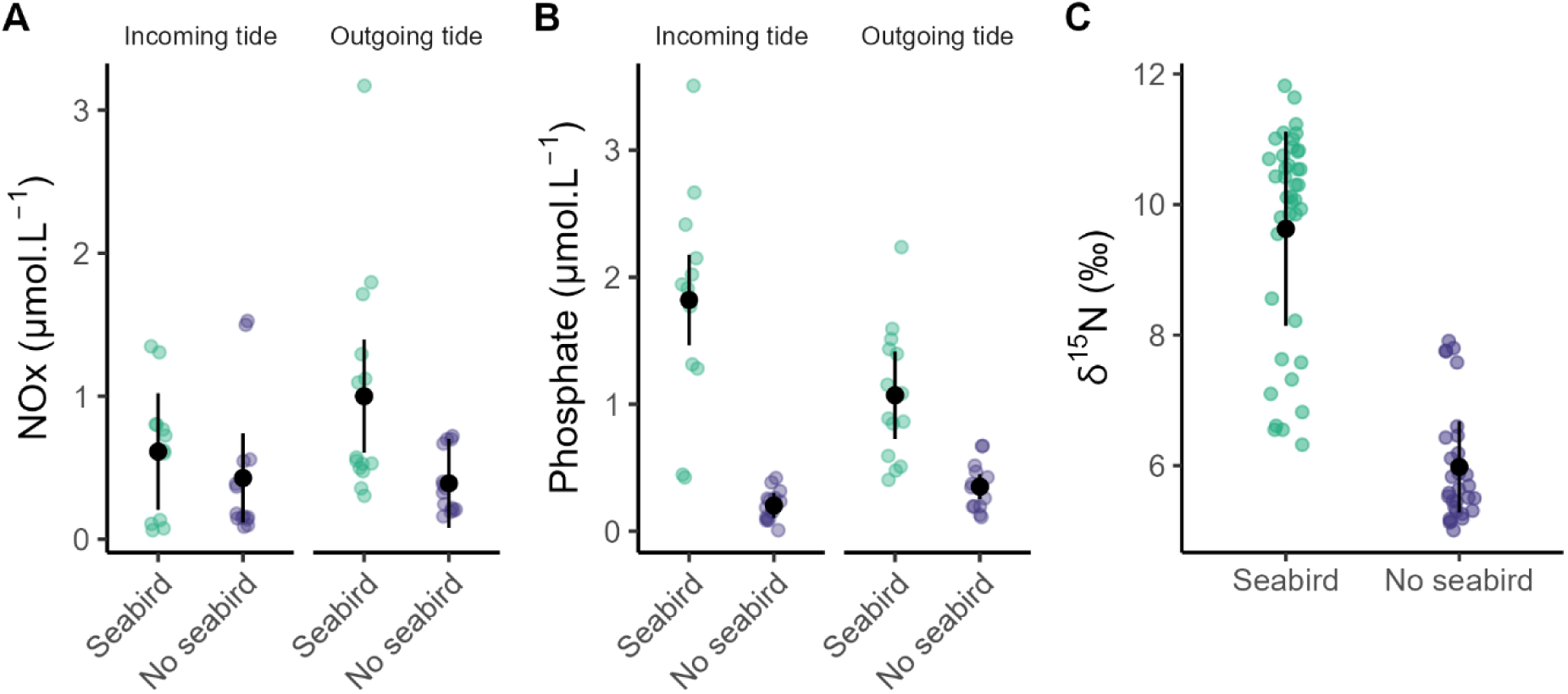
Nutrient levels in abiotic and biotic components adjacent to mangroves at seabird and non-seabird sites. Nutrient concentrations in surface seawater, NOx (A) and phosphate (B) during incoming and outgoing tides, and δ^15^N values of macroalgae *Halimeda* spp. (C) adjacent to mangroves, at seabird and non-seabird breeding sites on Aldabra Atoll. Black points and error bars display predicted means ± SD of linear mixed models and green and purple points display raw data. NOx = nitrate + nitrite.

## DISCUSSION

The ecological functions and ecosystem services provided by mangroves are reliant on the status and health of mangrove forests ^6^. It is therefore important to understand mangrove responses to natural processes that influence their status and health. Our study presents insights into the scale and extent of influence of cross-ecosystem nutrient subsidies on the ecology and function of Indo-Pacific mangroves and in the absence of human influence. Seabirds feed in open ocean and excrete substantial quantities of nutrients in mangrove forests where they nest. Mangroves with nesting seabirds were enriched in N and P and relieved of their nutrient limitations, although seabirds did not influence their nutrient resorption efficiencies. Through trophic pathways, seabird-derived nutrients were transferred to mangrove-associated invertebrate food webs. Furthermore, we documented the export of seabird-derived nutrients from mangroves to nearby coastal habitats through tidal flow, showing that the influence of nutrient contributions by mangrove-nesting seabirds extends beyond mangrove boundaries. Our study highlights the important roles of mobile consumers such as seabirds in maintaining functional connectivity between oceanic systems and coastal seascapes ^25^, and in improving the nutrient status of coastal habitats (Figure 4) and isolated island ecosystems. Our study adds more weight and nuance to the growing body of research on the influence of seabirds on highly productive coastal ecosystems ^26–28^.

**Figure 4.**
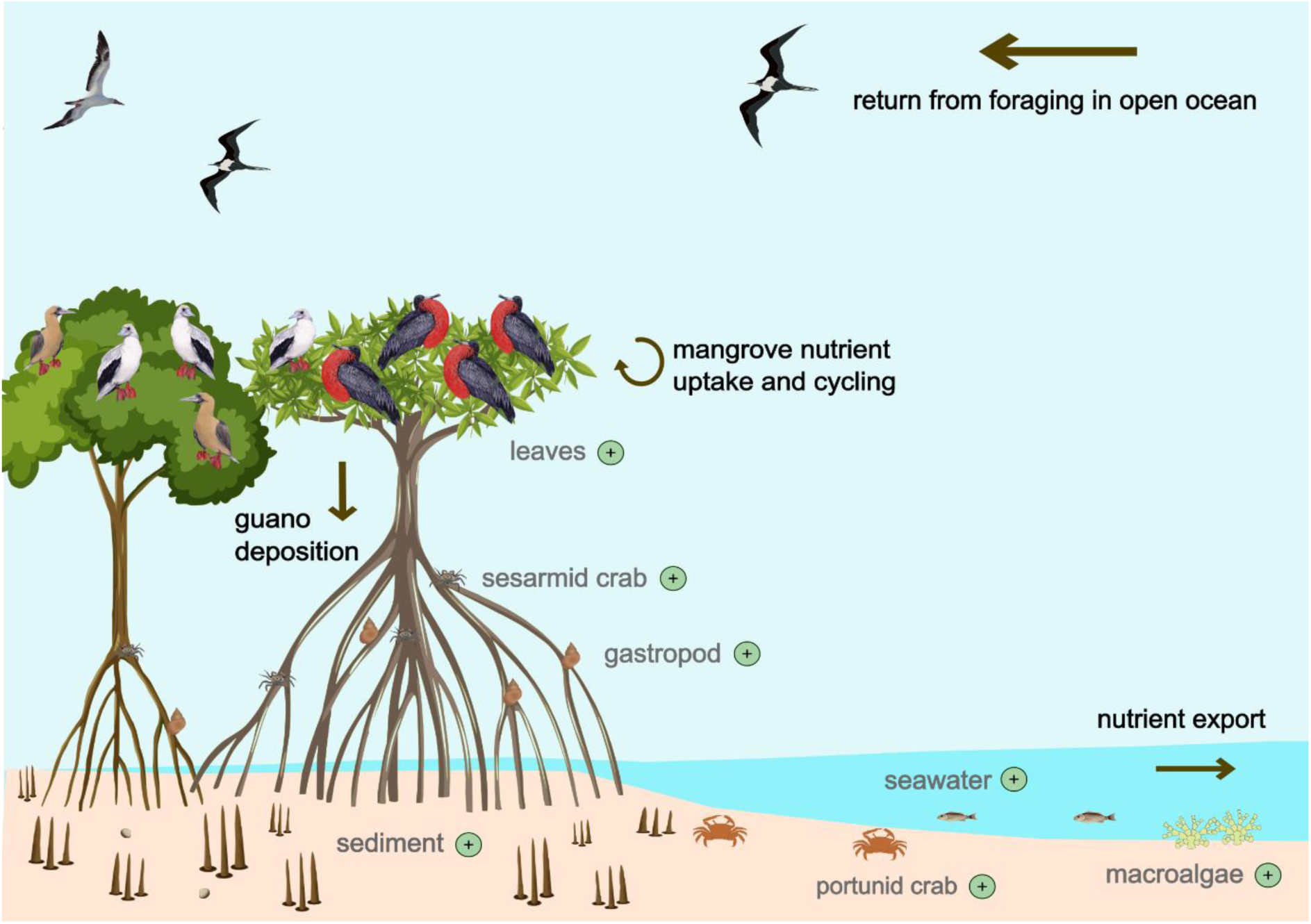
Summary figure illustrating the transfer of seabird-derived nutrients in mangrove forests. Seabirds forage at sea and deposit nutrient-rich guano in their mangrove breeding colonies. Seabird-derived nutrients enrich mangroves and associated invertebrate fauna, and are exported to adjacent habitats by tidal flow. Nutrient enrichment are indicated by plus (+) signs.

Oceanic atolls are particularly nutrient deficient as they are isolated from mainland nutrient sources and surrounded by oligotrophic waters. On atolls with seabird colonies, seabirds represent one of the main sources of marine-derived nutrient subsidies ^29^. Breeding red-footed boobies and frigatebirds delivered a combined total of 41.4 N tonne.year^−1^ and 40.9 P tonne.year^−1^ to Aldabra’s mangroves. This represents a minimum estimate since we did not account for non-breeding individuals. Furthermore, due to the bi-annual breeding of red-footed boobies and protracted breeding of frigatebirds ^30^, seabird-derived nutrient subsidies are delivered year-round. However, the majority of seabirds on Aldabra nest in tall mangrove trees along the northern and eastern lagoon shores ^24^. Accordingly, seabird nutrient contributions are disproportionately distributed, with Aldabra’s southern lagoon shores and terrestrial habitats deprived of seabird-derived nutrient subsidies. Additional sources of marine subsidies on Aldabra include algal wrack from seagrass beds ^31^, nesting sea turtles ^32^ and shorebirds ^33^, all of which contribute mainly to the sandy beach habitat around the atoll. Upwellings may also provide nutrient input to Aldabra’s coasts ^34^. Quantifying these additional sources and assessing their influence on the land- and seascape would provide a more holistic overview of marine nutrient budgets and functioning of marine-subsidized ecosystems on the atoll.

The mangrove forests in our study were dominated by *R. mucronata*, a species widespread in the Indo-Pacific region, and the only *Rhizophora* species occurring in the Western Indian Ocean ^35,36^. The higher δ^15^N concentration found in sediment and leaves at seabird sites confirms that seabird-derived nutrients are transferred to mangroves. Foliar N and P concentrations of *R. mucronata* were 39% and 28% higher, respectively, at seabird sites, providing evidence of seabird-derived nutrient uptake by mangroves. Our results corroborate documented increases in nutrients at seabird sites relative to control sites for coastal *Rhizophora* species in Central and North America, for example *R. mangle* (Belize: +37% N ^37^; Honduras: +41% N and +20% P ^18^; Mexico: +25 %N and +17 %P ^16^; Florida: +33 %N ^15^). Mangroves on Aldabra grow in a lagoonal low-lying carbonate environment and do not receive large volumes of terrigenous sediment supply and nutrients compared to mangroves growing in deltas or estuaries ^38,39^. In this nutrient-limiting environment, our results show that seabirds contribute essential macronutrients and improve the nutrient status of mangroves.

Responses of mangroves to nutrient enrichment are site- and species-specific, with the main limiting nutrient of the site governing species nutrient requirement and tolerances ^40^. Nutrient limitation in mangroves is usually attributed to N or P limitation ^41^. Foliar N:P ratios can be used to infer nutrient limitation ^42^, with N:P ratios > 32 indicating P limitation in mangroves ^7,43^. Foliar N:P ratio in our study was 11.7 ± 2.45 (Mean ± SD), suggesting mangroves in our study are N-limited, which is typical for fringe mangroves influenced by frequent tidal flushing ^7,44^. N limitation at our study site is further indicated by the equally low N:P ratios documented in previous analyses on mangrove soil ^45^ and lagoonal sediment porewater on Aldabra ^46^. At our seabird sites, uptake of seabird-derived nutrients alleviated mangrove nutrient limitations of both N and P, observed by reductions in foliar C:N and C:P ratios of *R. mucronata*. Nutrient enrichment in N-limited mangroves can generate multiple higher-order effects. For example, in Florida, seabird-derived nutrients increased primary production, leaf nutritional value and herbivory ^15^. Similarly, experimental nutrient enrichment in N-limited mangroves has demonstrated increases in shoot biomass and tree growth ^40^.

Resorption of nutrients prior to leaf fall, a process in which nutrients are resorbed from senescent leaves and are directly available for continued plant growth, is a vital nutrient recycling and conservation mechanism in mangroves ^7,47^. When nutrients become available, mangroves reduce their nutrient resorption efficiency ^48,49^. Based on this, we expected reductions in resorption efficiency at our seabird sites compared to non-seabird sites; however, no differences in RE-N or RE-P were detected. Furthermore, given that our study site is N-limited, we would expect greater resorption of N compared to P ^47^, however, average RE-N values were similar to average RE-P values at both seabird and non-seabird sites. Seabird-derived nutrient enrichment has been documented to reduce mangrove resorption efficiency in Mexico ^16^ but not in Belize ^37^. Similar contrasting patterns are shown in experimental nutrient enrichment studies ^50,51^, suggesting internal nutrient cycling by mangroves is complex, regulated by the interacting effects between species-specific physiological capacity to conserve nutrients ^49^, nutrient demand (through increased growth rate) and supply ^7,51^.

A consequence of nutrient enrichment is that it can reduce stability and resilience of mangrove forests ^9,52,53^. Nutrient enrichment causes plants to invest more in aboveground growth and less in belowground biomass ^54,55^. Furthermore, nutrient availability increases rates of microbial decomposition of organic matter ^48,56^. A forest with lower root biomass and higher decomposition rates, is likely to have reduced accretion of sediment organic matter ^57^, resulting in reduced shoreline stability ^52^ and increased probability of mangrove death when faced with environmental stressors or events ^9,53^. On this basis, seabird-derived nutrient enrichment has been linked to declines of mangrove cays in Belize ^37^. However, in contrast to Belize, some of the greatest increases in mangrove extent on Aldabra over the past two decades coincided with seabird nesting locations ^58^. Furthermore, mangroves on Aldabra are far from urban or agricultural centers compared to Belize, and are therefore not influenced by additional anthropogenic nutrient sources. Natural nutrient sources provide N and P in optimal ratios ^59,60^, generating contrasting responses in coastal habitat structure and functions compared to anthropogenic sources ^61^. In addition, seabirds on Aldabra nest primarily in tall trees in the mangrove fringes, which receive constant tidal flushing, limiting over-enrichment of nutrients ^9^. Altogether, this suggests that seabird-derived nutrient enrichment is unlikely to be causing declines in mangroves on Aldabra. Soil nutrient content is a main driver of mangrove aboveground biomass on Aldabra ^45^. Therefore, by fertilizing mangroves where they nest, seabirds likely enhance mangrove productivity in their breeding areas, further promoting suitable nesting habitat, likely creating a positive-feedback loop for seabird populations ^10^. Indeed, both mangroves and mangrove-nesting seabird populations on Aldabra have increased over the last few decades ^23,58^.

Multiple pathways of energy and nutrient flow exist in mangrove food webs due to the wide range of available food sources, such as plankton, benthic microalgae, mangroves, macroalgae and macrophytes ^4,62^. By assessing the trophic transfer of seabird-derived nutrients, we found that seabirds also enriched the mangrove invertebrate food web. We detected higher δ^15^N at seabird sites at all trophic levels, including baseline (sediments), primary producers (mangrove leaves), primary consumers (gastropods and sesarmid crabs) and a secondary consumer (portunid crab). Invertebrate food web enrichment by seabird guano has been documented in other coastal ecosystems such as littoral habitats ^63^, fjords ^64^ and coastal ponds ^65^, but not previously in mangrove forests. Invertebrates are key components of mangrove-associated macrofauna, playing substantial roles in nutrient cycling of detritus ^7^. Further research should explore additional pathways of seabird-derived nutrient flow in mangroves, as well as how seabird nutrient enrichment affects invertebrate trophic structure, interactions and ecological functions.

Given the absence of anthropogenic influence in Aldabra’s lagoon, the higher levels of NOx and phosphate in seawater adjacent to mangroves with seabird colonies can be attributed to seabird guano inputs. Increased levels of seawater nutrients around seabird colonies in remote oligotrophic locations confirms seabird-derived nutrients as an important source of marine productivity ^66^. Indeed, δ^15^N levels of macroalgae *Halimeda* spp. growing adjacent to mangroves were also higher compared to non-seabird sites. These results indicate horizontal transfer of seabird-derived nutrients, from mangroves through tidally-mediated nutrient exchange. Nutrients can also be transferred to adjacent coastal habitats (such as coral reefs or seagrass areas) via mangrove leaf litter, which plays a key role in sustaining adjacent marine food webs ^62^. Leaves with higher-nutrient content or nutritive value are more rapidly broken down than less nutrient-rich leaves ^15^. Seabird-derived nutrients from mangroves may therefore extend to adjacent marine communities through trophic pathways, for example via fish communities from nearby reefs or seagrass beds that utilize mangroves as nursery or feeding habitat.

Mangroves on Aldabra support a wide range of biodiversity and ecosystem service benefits. In addition to supporting large breeding populations of seabirds, Aldabra’s mangroves sustain threatened and regionally important populations of numerous marine species. Aldabra’s mangroves provide nursery habitat for one of the largest green turtle populations in the region ^32^, play a critical role as feeding, breeding and nursery habitats for numerous bony fish and shark species ^67^. Aldabra’s mangroves support the highest biomass of herbivorous fish and the highest abundance of sharks in Seychelles ^68,69^. Furthermore, Aldabra’s mangroves comprise one of Seychelles’ largest blue carbon ecosystems ^45^. By improving mangrove nutrient status and health, we show that seabirds nesting on Aldabra boost biodiversity and ecosystem services.

Our study provides important insights to nutrient contribution, cycling and transfer in a system with strong land-sea connectivity and without human stressors. We present critical data to start unravelling the mechanisms and extent of nutrient connectivity in mangroves, linked to the cross-ecosystem ocean-derived subsidies provided by seabirds to islands. Given the critical roles of mobile consumers such as seabirds in maintaining ecosystem health and functions, conservation and management actions should focus on restoring seabird populations and their breeding grounds. Such efforts will present maximum benefits for people and biodiversity at multiple scales ^70^.

## EXPERIMENTAL PROCEDURES

### Resource availability

#### Lead contact

Further information and requests for resources should be directed to and will be fulfilled by the lead contact, Jennifer Appoo

### Materials availability

This study did not generate new unique materials.

### Data and code availability

Data and code to replicate the findings presented will be made available on Dryad Digital Repository upon acceptance.

Any additional information required for reanalyzing the data reported in this paper is available from the lead contact upon request.

### Study site

Aldabra Atoll (9°24′ S, 46°20′ E) is part of the Seychelles archipelago (Figure S1). The atoll comprises four main islands (land area 15,500 ha, mean elevation 8 m asl), encircling a large shallow lagoon (20,500 ha) and separated by deep (10‒30 m depth) narrow channels ^71,72^. Aldabra’s mangroves occur primarily along the lagoon shores, covering 1,720 ha ^73^. Mangrove habitat on the northern and south-eastern lagoon shores consists of tall trees forming closed canopies, while mangrove areas on Aldabra’s southern shores are smaller and scattered (see Table S1). Seven mangrove species occur, with *Rhizophora mucronata*, *Bruguiera gymnorhiza*, *Ceriops tagal* and *Avicennia marina* being the most common ^74^. Mangrove sediments are mainly sand (53%) and loamy sand (26%) and with an average salinity of 8.3 ± 3.1 (practical salinity scale) ^58^. Aldabra is influenced by a semi-diurnal meso-tidal regime (range: 2–3 m), resulting in strong tidal currents that drain approximately 75% of the lagoon at low tide through the main channels ^72,75^. Rainfall patterns on Aldabra show marked seasonal variations influenced by monsoon winds. The majority of rain is concentrated between November and April with a mean of 975 mm.year^−1^, however rainfall does not influence primary productivity of the mangroves ^76^.

On Aldabra, great and lesser frigatebirds, *Fregata minor* and *F. ariel* respectively, nest together in four separate colonies, with a combined breeding population estimated at 16,534 pairs (SIF, *unpubl. data*). The red-footed booby *Sula sula* nests together with or outside the frigatebird colonies, totaling 36,720 pairs (SIF, *unpubl. data*). Great frigatebird breeding peaks in October, while lesser frigatebirds and red-footed boobies have two breeding peaks, in June and October, and February and August, respectively (SIF, *unpubl. data*). The distribution of mangrove-nesting seabirds is not uniform, with seabirds found only in mangroves on the northern, eastern and south-eastern lagoon shores ^30^. Mangroves and seabirds on Aldabra are recognized globally, as a Ramsar Wetland Site of International Importance and an Important Bird Area (BirdLife International), respectively. Furthermore, Aldabra’s mangroves and seabirds have been strictly protected since designation of the atoll as a Strict Nature Reserve in 1981 and UNESCO World Heritage Site in 1982.

### Experimental design

We investigated the effects of breeding seabirds on Aldabra’s mangroves during the nesting period of all three species (November 2022 to March 2023). Sampling was conducted at 10 sites comprising five sites with no or low numbers of nesting seabirds (< 20 nests) and five sites with high numbers of nesting seabirds (Figure S1). Sites were distributed along the lagoon shore around the atoll and situated at least 1 km apart. All sampling sites were located on the seaward fringe of mangroves, which is where the majority of seabirds roost and nests on Aldabra. Within each site, we conducted mangrove measurements within eight quadrats of 5 x 5 m, separated by a minimum of 50 m. Mangrove sampling was conducted at spring low tide when mangroves and intertidal areas were exposed.

### Seabird nutrient contributions

To determine the amount of nutrients deposited by mangrove-nesting seabirds, we collected droppings around nests of red-footed boobies and frigatebirds. For the latter, droppings were sampled around nests with big chicks. Given the difficulty of correctly identifying frigatebird species with chicks, the samples were generalized to represent both great and lesser frigatebirds. Droppings were kept cool in the field and then stored frozen in the laboratory until further processing. Total nitrogen (N) content in seabird droppings was obtained by mineralization according to the Kjeldahl method ^77^ (Büchi KjelMaster K-375, Switzerland). Total phosphorus (P) was determined by preliminary dry combustion (600 °C), then dissolving the residual material in 6 M HCl. P concentration was obtained by extraction using ammonium molybdate and ascorbic acid and measured with a continuous flow colorimeter (Proxima, Alliance Instrument, USA). Nutrient analyses were conducted at the CIRAD laboratory of agronomy in Reunion Island (France). Additional seabird droppings were collected for isotope analysis (see below).

We used N and P concentrations in seabird droppings to estimate the total annual N and P input from breeding seabirds using previously used methods ^78,12^:

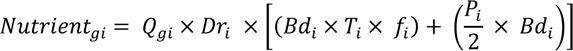

where *Nutrient*_*gi*_, t.yr^−1^ is the annual input per nutrient type and species, *Q*_*gi*_, mg.g^−1^ is the concentration of N or P measured in seabird droppings for each species, *Bd*_*i*_ is number of breeding adults for that species, *T*_*i*_ is the length of the breeding period (number of days from courtship to chick-rearing, *f*_*i*_ is the proportion of time spent at the colony during breeding to account for absence due to feeding forays and *P*_*i*_ is the productivity of the species (fledged chicks per breeding pair). Defecation rate for red-footed boobies was obtained from Young et al., ^79^. For frigatebirds, we estimated defecation rates based on red-footed booby measurements, scaled allometrically to body size following Staunton-Smith and Johnson ^80^, and using a median value of the adult body sizes of lesser and great frigatebirds. Values for the length of the breeding period, proportion of time at the colony, productivity and adult body sizes for each species were obtained from Riddick et al., ^78^. Our estimates do not account for nutrient contributions by non-breeding individuals since we lacked data on numbers and diurnal movement patterns.

### Mangrove forest structure

We assessed forest characteristics in each quadrat following methods previously employed on Aldabra ^45^. We recorded the species of all trees >2 m height and measured their diameter at breast height (DBH, 1.3 m height) using a measuring tape, following guidelines for trees with anomalies such as prop roots ^81^. We counted the number of seedlings and used an inclinometer (Suunto PM-5/66 PC, Vantaa, Finland) to measure tree height to the nearest ± 0.5 m. We applied the allometric equation by Chave et al., ^82^ to determine aboveground biomass (ABG) following Constance et al., ^45^. Aboveground biomass (ABG, kg) per tree was obtained from the wood density (*ρ*, g.cm^−3^), the DBH (D, cm), and height (H, m).

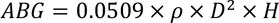

### Mangrove foliar nutrient analyses

To test the effects of seabird subsidies on mangrove nutrients, we measured leaf nutrient concentrations of the dominant species *R. mucronata*. In each sampling quadrat, we collected six young fully expanded green leaves exposed to the sun and six yellow (senescent) leaves from at least three individual trees. The leaves were pooled per quadrat to form one composite green and yellow leaf sample. Leaves were rinsed with fresh water, oven-dried at 60°C for a minimum of 48 hrs and powdered using a ball mill. Total carbon (C) and N were determined using an elemental-analyser (Elementar Vario Micro Cube, Lancaster University, UK). Total P content was determined by the Olsen-Dabin method using 0.5 M sodium hydrogen carbonate solution with a continuous flow colorimeter (Proxima, Alliance Instrument, USA) at the CIRAD laboratory of agronomy in Reunion Island (France). Sub-samples were obtained for isotope analysis.

We derived foliar nutrient ratios (C:N, C:P mass basis) to explore nutrient limitations to mangrove growth ^7,16^. As an indicator of nutrient cycling, we calculated the nutrient resorption efficiency of N (RE-N) and P (RE-P), which corresponds to the percentage of N or P recovered from senescing leaves prior to leaf fall ^40,47^, based on the equation by Chapin and Cleve ^83^:

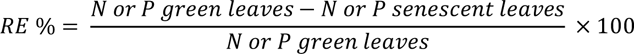

### Isotope sampling and analyses

To assess the transfer of seabird-derived nutrients in mangroves we conducted isotopic analyses on seabird droppings, mangrove sediment and green *R. mucronata* leaves. Surface sediments were sampled by inserting a PVC corer (⌀ = 22 mm) in the top 2 cm sediment layer at three random positions in each quadrat. To assess the trophic transfer of seabird-derived nutrients to mangrove fauna we conducted isotopic analyses on mangrove invertebrates comprising of gastropods and crabs occupying different trophic levels. As primary consumers, we collected herbivorous gastropods of the genus *Littoraria* from mangrove roots and trunk within each quadrat, and leaf-eating sesarmid crab *Sesarma leptosoma* on mangrove trees at each site. As secondary consumers, we collected omnivorous portunid crab *Thalamita crenata* on the intertidal mudflat adjacent to mangrove forests. We sampled foot muscle from gastropods, body muscle from sesarmid crabs and limb muscle from portunid crabs. We collected macroalgae on rocky outcrops of the intertidal mudflat and within 50 m of the mangrove fringe for isotopic analyses to assess nutrient uptake in the adjacent habitat. We selected macroalgae since they incorporate nutrients over a relatively long period (weeks to months), constituting an ideal proxy for ambient nutrient conditions ^84^. At each site, we randomly collected thalli of the macroalgae *Halimeda* spp. The latter has proven a good proxy for seabird nutrients since it responds positively to seabird subsidies ^60^.

Isotopic values of seabird-derived nutrients are enriched in nitrogen (δ^15^N) due to the high trophic position of seabirds in the marine food chain and enrichment of δ^15^N in seabird guano after deposition ^85^. δ^15^N is therefore used to trace seabird-derived nutrients in recipient communities ^27^. All samples for isotopic analyses were dried at 60°C and ground into a homogeneous powder using a ball mill. Isotopic ratios were measured using an Isoprime 100 Isotope Ratio Mass Spectrometer with international standards IAEA 600, USGS 41 and CH6, at the stable isotope facility at Lancaster University (Lancaster, UK). Accuracy based on internal standards was within 0.1 ‰ SD and selected samples were run in triplicate to further ensure accuracy of readings.

### Surface seawater nutrient concentrations

The semi-diurnal tidal fluctuations at Aldabra generate large and rapid changes in water levels, causing strong tidal currents ^72^. Most of the guano inputs by mangrove-nesting seabirds on Aldabra are deposited directly in the water or are washed away during the subsequent high tide when deposited at low tide ^86^. To assess nutrient export from the seabird colonies, we sampled surface seawater in 30-ml containers at each site on two occasions; on the incoming tide while the lagoon fills and on the outgoing tide while the lagoon drains. Samples were collected in duplicate at each location and occasion and treated at 80°C in a drying oven for a minimum of 2.5 hrs. Samples were stored at room temperature in the dark until laboratory analysis. NOx (nitrate NO ^−^ + nitrite NO ^−^) and phosphate (PO_4_ ^3−^) concentrations were determined using an AA3 auto-analyser (Seal Analytical) following the method of Aminot and Kérouel ^87^. Samples were processed at IRD laboratory in Plouzané (France) and had an accuracy of 0.5 μmol.L^−1^ and 0.7 μmol.L^−1^, for NOx and PO ^3−^, respectively.

### Data analyses

We assessed the influence of seabirds on nutrient parameters using univariate tests of differences between seabird and non-seabird sites. We formulated linear mixed models (LMMs) for each nutrient parameter (%N, %P, C:N, C:P, RE-N, RE-P) of mangrove leaves and δ^15^N for mangrove leaves, sediment, gastropod, crabs and macroalgae with seabird status (seabird/no seabird) as fixed effect and site as random effect. For some parameters, variances were heterogeneous between the predictor levels (i.e., higher within seabird than non-seabird sites), therefore we included the ‘VarIdent’ error structure on seabird status, which allows different variance of predictor levels. We assessed residual spatial autocorrelation of each model by plotting residuals against location and found no spatial autocorrelation in any of these models. This was confirmed by comparing models with and without spatial weights matrix using Akaike’s Information Criterion (AIC), and models without had the lowest AIC values. Model diagnostics were performed by plotting residuals against fitted values and explanatory variables to verify assumptions of homogeneity of variance and independence following Zuur and Ieno ^88^.

We assessed differences in surface seawater nutrient concentrations separately for NOx and phosphate. We used LMMs with seabird status, including an interaction with tidal regime (incoming/ outgoing) as fixed effects and site as random effect. These models included the ‘VarIdent’ error structure on seabird status. Where significant differences were detected, we performed post-hoc tests using the *emmeans* package ^89^. Model residuals were assessed to verify model assumptions and to confirm there were no spatial dependencies ^88^. All models were formulated using the function ‘lme’ from package *nlme* ^90^ and analyses were conducted on R version 4.3.0 ^91^.

## SUPPLEMENTAL INFORMATION

Document S1: Figure S1 and Tables S1-S3.

Supplemental Information includes one figure and three tables and can be found online at:

## ACKNOWLEDGMENTS

We thank Seychelles Islands Foundation staff at head office and on Aldabra Atoll for providing administrative and logistical support during the fieldwork. Funding was provided by the African World Heritage Fund through a Moses Mapesa Research grant to J.A, the PADI Foundation marine research grant to J.A, and the Bertarelli Foundation under the Bertarelli Marine Science Program to N.A.J.G. J.A was supported by a doctoral fellowship from the Reunion Island Regional Council. Fieldwork was conducted with approval of the Seychelles Bureau of Standards under permit number A0157 and the Seychelles Islands Foundation. Icons for Figure 4 were obtained from University of Maryland Center for Environmental Science, Integration and Application Network (https://ian.umces.edu/media-library/), Freepik (https://www.freepik.com/) and Birds Caribbean (https://www.birdscaribbean.org/).

## AUTHOR CONTRIBUTIONS

Conceptualization: J.A, N.B, N.A.J.G and S.J; Methodology: J.A and N.B; Data collection: J.A; Formal analysis: J.A; Writing – Original Draft: J.A; Writing – Review and Editing: N.A.J.G, N.B, S.J, A.L and M.L.C.

## DECLARATION OF INTERETS

The authors declare no competing interest.

