## Supplementary material for "Seabird nutrient subsidies enrich mangrove ecosystems and are exported to nearby coastal habitats": Document S1: Figure S1 and Tables S1-S3.

**SUPPLEMENTAL INFORMATION**

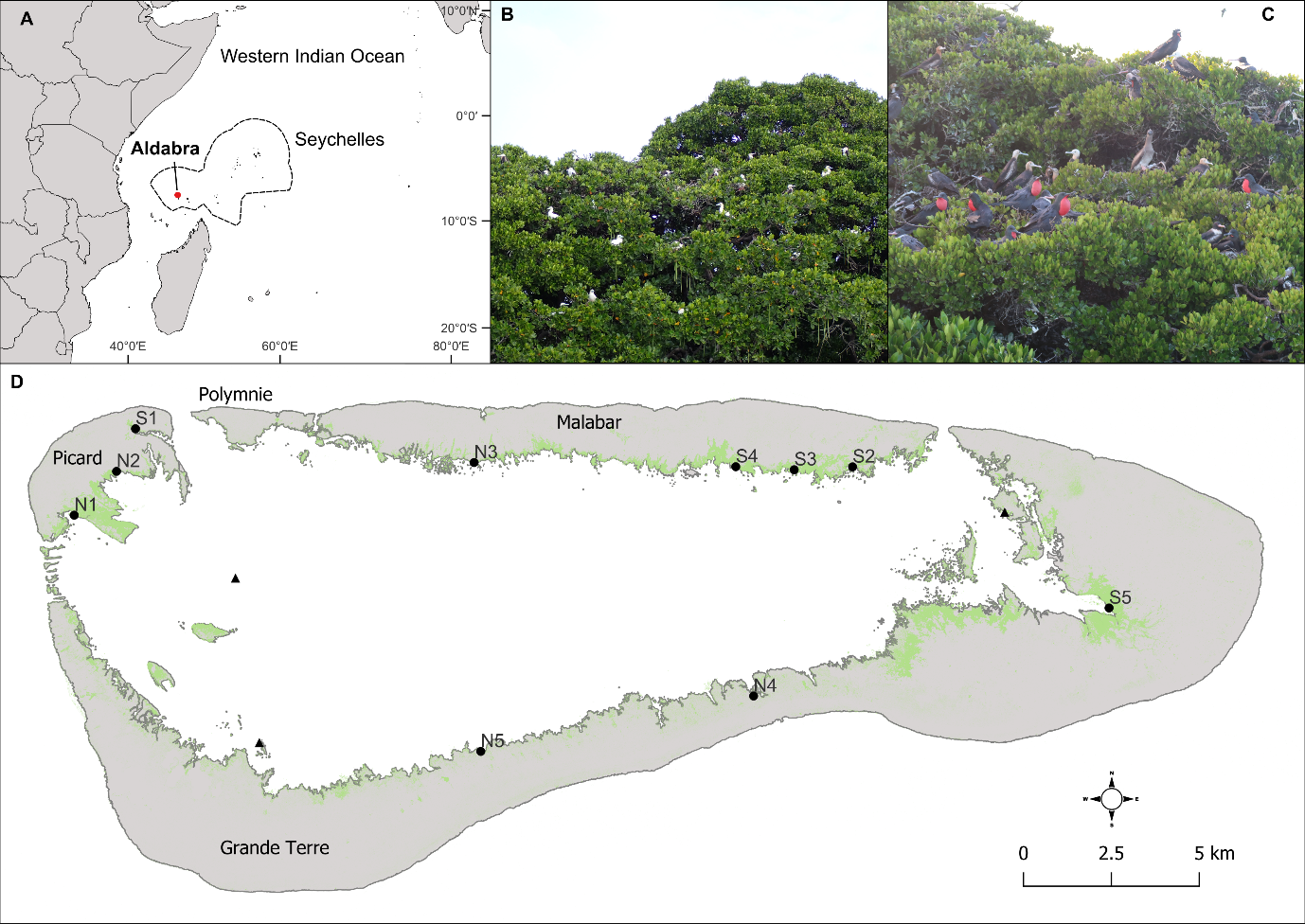

**Figure S1. Study site and sampling locations.**

(A) Location of Aldabra Atoll (Seychelles) in the Western Indian Ocean.

(B) Red-footed boobies nesting in mangroves at Aldabra.

(C) Frigatebird breeding colony in mangroves at Aldabra.

(D) Sampling locations (circles) on Aldabra at sites with seabirds (S1–S5) and sites with no/few seabirds (N1–N5), with mangrove distribution shown in green. Triangles show locations of additional seawater sampling sites in the lagoon.

**Table S1. Mangrove forest structure at each site on Aldabra Atoll.**

Sites with nesting seabirds (S1–S5) and no nesting seabirds (N1–N5). Community composition of main mangrove species shown; Rm: *Rhizophora mucronata*, Bg: *Brugueira gymnorrhiza*, Ct: *Ceriops tagal*. DBH: diameter at breast height.

| Site | Species composition | | | Tree density (trees. ha^-1^) | Tree DBH (cm) | Tree height (m) | Above-ground biomass (Mg. ha^-1^) |
| --- | --- | --- | --- | --- | --- | --- | --- |
|  | % Rm | % Bg | % Ct | ̶̶ ̶̶ ̶̶ ̶̶ ̶̶ ̶̶ ̶̶ ̶̶ ̶̶ ̶̶ ̶̶ ̶̶ ̶̶ ̶̶ ̶̶ ̶̶ ̶̶ Mean ± SD ̶̶ ̶̶ ̶̶ ̶̶ ̶̶ ̶̶ ̶̶ ̶̶ ̶̶ ̶̶ ̶̶ ̶̶ ̶̶ ̶̶ ̶̶ ̶̶ | | | |
| Seabird | |  |  |  |  |  |  |
| S1 | 100 | 0 | 0 | 2700 ± 1885 | 12.6 ± 6.6 | 7.4 ± 2.5 | 219.5 ± 246.6 |
| S2 | 88 | 9 | 3 | 4650 ± 1788 | 13.3 ± 5.5 | 7.2 ± 2.1 | 335.8 ± 114.7 |
| S3 | 84 | 10 | 6 | 4100 ± 1957 | 10.1 ± 5.1 | 6.9 ± 2.4 | 195.4 ± 84.6 |
| S4 | 65 | 26 | 9 | 4600 ± 1750 | 10.9 ± 5.8 | 7.5 ± 2.9 | 254.4 ± 126.2 |
| S5 | 98 | 0 | 0 | 4350 ± 4957 | 8.1 ± 5.8 | 6.9 ± 3.2 | 178.1 ± 105.6 |
| No seabird | | |  |  |  |  |  |
| N1 | 78 | 13 | 9 | 9450 ± 4017 | 6.3 ± 4.2 | 4.4 ± 2.1 | 160.3 ± 90.2 |
| N2 | 99 | 0 | 1 | 5150 ± 2708 | 9.3 ± 4.4 | 6.5 ± 2.3 | 194.8 ± 83.8 |
| N3 | 52 | 8 | 40 | 9350 ± 6552 | 8.3 ± 5.3 | 4.8 ± 2.3 | 252.5 ± 134.5 |
| N4 | 96 | 0 | 4 | 1350 ± 1299 | 5.6 ± 2.9 | 2.2 ± 0.2 | 7.1 ± 6.48 |
| N5 | 78 | 22 | 0 | 450 ± 334 | 5.7 ± 4.6 | 2.3 ± 0.2 | 3.1 ± 3.3 |

**Table S2.** **Results of linear mixed models to test differences of nutrient parameters between seabird and non-seabird sites.**

Significant differences (*P* ≤ 0.05) are marked in bold. RE-N: nitrogen resorption efficiency, RE-P: phosphorus resorption efficiency.

Related to Figure 1, 2 and 3.

| Nutrient parameter | F-value | *P*-value | Conditional R2 | Marginal R2 |
| --- | --- | --- | --- | --- |
| Mangrove leaves *Rhizophora mucronata* (N_obs_ = 80, N_site_ = 10) | | | |  |
| % N | 50.7 | **< 0.0001** | 0.75 | 0.71 |
| % P | 10.2 | **0.013** | 0.61 | 0.38 |
| C:N | 44.1 | **< 0.0001** | 0.51 | 0.47 |
| C:P | 6.86 | **0.031** | 0.40 | 0.19 |
| RE-N % | 2.61 | 0.15 | 0.40 | 0.11 |
| RE-P % | 1.59 | 0.24 | 0.53 | 0.09 |
| δ^15^N ‰ | 5.09 | **0.05** | 0.66 | 0.24 |
| Sediment (N_obs_ = 240, N_site_ = 10) | |  |  |  |
| δ^15^N ‰ | 21.4 | **0.0017** | 0.95 | 0.65 |
| Gastropod (N_obs_ = 240, N_site_ = 10) | |  |  |  |
| *Littoraria* spp. δ^15^N ‰ | 113.8 | **< 0.0001** | 0.68 | 0.64 |
| Sesarmid crab (N_obs_ = 150, N_site_ = 10) | |  |  |  |
| *Sesarma leptosoma* δ^15^N ‰ | 35.2 | **< 0.001** | 0.93 | 0.73 |
| Portunid crab (N_obs_ = 86, N_site_ = 9) | |  |  |  |
| *Thalamita crenata* δ^15^N ‰ | 16.8 | **0.005** | 0.83 | 0.55 |
| Macroalgae (N_obs_ = 70, N_site_ = 7) | | |  |  |
| *Halimeda* spp. δ^15^N ‰ | 11.0 | **0.021** | 0.94 | 0.58 |

**Table S3. Results of linear mixed models to test differences in surface seawater nutrients.**

Significant differences (*P* ≤ 0.05) are marked in bold.

Related to Figure 3.

|  | Fixed effects | F-value | P-value | Conditional R2 | Marginal R2 |
| --- | --- | --- | --- | --- | --- |
| Lagoon surface seawater nutrients | |  |  |  |  |
| **NOx** | Seabird | 2.80 | 0.12 | 0.85 | 0.23 |
| (N_obs_ = 54, N_site_ = 14) | Tide | 0.22 | 0.64 |  |  |
|  | Seabird: Tide | 5.76 | **0.021** |  |  |
| **Phosphate** | Seabird | 34.1 | **0.0001** | 0.97 | 0.78 |
| (N_obs_ = 54, N_site_ = 14) | Tide | 4.02 | **0.052** |  |  |
|  | Seabird: Tide | 17.8 | **0.0001** |  |  |
